# The chemistry and pharmacology of putative synthetic cannabinoid receptor agonist (SCRA) new psychoactive substances (NPS) 5F-PY-PICA, 5F-PY-PINACA, and their analogues

**DOI:** 10.1101/430959

**Authors:** Samuel D. Banister, Richard C. Kevin, Lewis Martin, Axel Adams, Christa Macdonald, Jamie J. Manning, Rochelle Boyd, Michael Cunningham, Marc Y. Stevens, Iain S. McGregor, Michelle Glass, Mark Connor, Roy R. Gerona

## Abstract

The structural diversity of synthetic cannabinoid receptor agonist (SCRA) new psychoactive substances (NPS) has increased since the first examples were reported a decade ago. 5F-PY-PICA and 5F-PY-PINACA were identified in 2015 as putative SCRA NPS, although nothing is known of their pharmacology. 5F-PY-PICA, 5F-PY-PINACA, and analogues intended to explore structure-activity relationships within this class of SCRAs were synthesized and characterized by nuclear magnetic resonance spectroscopy and liquid chromatography–quadrupole time-of-flight–mass spectrometry. Using competitive binding experiments and fluorescence-based plate reader membrane potential assays, the affinities and activities of all analogues at cannabinoid type 1 and type 2 receptors (CB_1_ and CB_2_) were evaluated. All ligands showed minimal affinity for CB_1_ (p*K*_i_ < 5), although several demonstrated moderate CB_2_ binding (p*K*_i_ = 5.45–6.99). At 10 μM none of the compounds produced an effect > 50% of CP55,950 at CB_1_, while several compounds showed a slightly higher relative efficacy at CB_2_. Unlike other SCRA NPS, 5F-PYPICA and 5F-PY-PINACA did not produce cannabimimetic effects in mice at doses up to 10 mg/kg.

## Introduction

More than 800 new psychoactive substances (NPS) have been identified in the past decade, including more than 250 synthetic cannabinoid receptor agonists (SCRAs).^1,2^ The earliest SCRA NPS were 1-substituted 3-acylindoles, like JWH-018 (**1**, Fig. 1), and were based on research compounds with activity at cannabinoid type 1 and 2 (CB_1_ and CB_2_, respectively) receptors disclosed by academic laboratories or pharmaceutical companies.^3–5^ As acylindole SCRAs were prohibited in various jurisdictions, newer derivatives not yet subject to legislation such as AB-FUBINACA (**2**) and MDMB-CHMICA (**3**) appeared to replace them.^6,7^ SCRAs represent the largest and most chemically heterogeneous class of NPS, and new examples continue to emerge in drug markets around the world.^8–13^

Unlike early SCRA NPS, many recent SCRAs have no precedent in the scientific literature when first detected in drug markets, necessitating robust analytical workflows for identification.^14^ Even for the most well-studied SCRA NPS, existing data pertain to the activity of these compounds in cellular systems or animal models,^15–18^ with very few human studies reported.^19^ Of greater concern is the trend for increasing potency of SCRA NPS, with the most recent examples frequently associated with serious adverse effects including myocardial infarction, ischemic stroke, acute kidney injury, tonic-clonic convulsion, coma, and death.^20–30^

**Figure 1.**
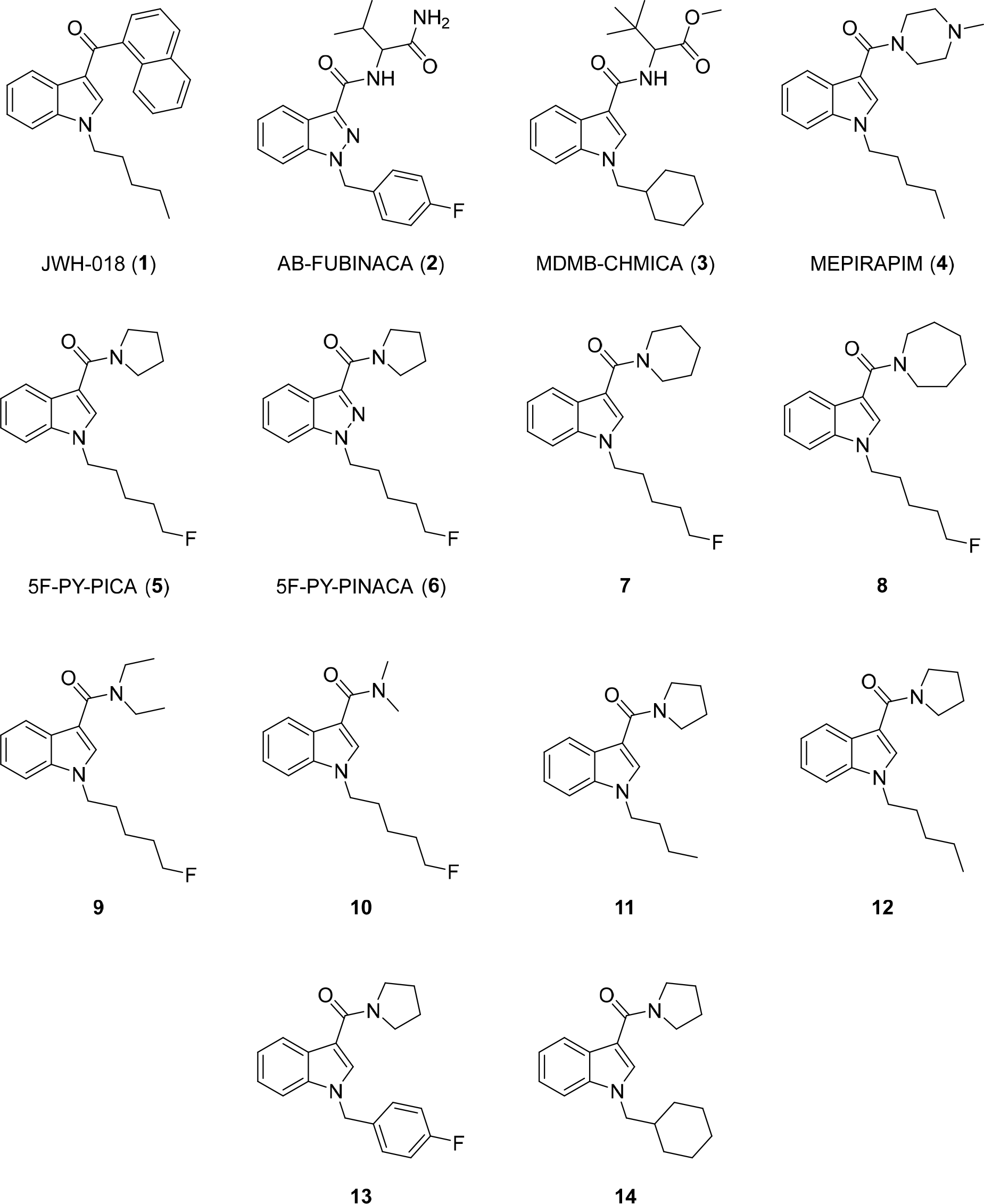
Selected synthetic cannabinoid receptor agonist (SCRA) new psychoactive substances (NPS) and their prophetic analogues.

The largest class of SCRA NPS are 1-substituted indole- and indazole-3-carboxamides. These typically feature secondary amides, as found in **2** and **3**, however, tertiary amides comprised of alicyclic or heterocyclic substituents have occasionally been reported. The earliest example was piperazine derivative MEPIRAPIM (**4**), reported in Japan in 2014,^31^ likely inspired by a class of water-soluble cannabinoid analgesics developed by Schering-Plough.^32^ The first putative SCRA NPS featuring a pyrrolidine ring at the 3-position were recently reported; indole 5F-PY-PICA (**5**) was identified in France in June 2015,^33^ and the indazole analogue 5F-PY-PINACA (**6**) was notified in Sweden around the same time.^33^ The metabolic profile of 5F-PY-PICA in rat and human hepatocytes was recently described, but nothing else is known of the pharmacology of this class of putative SCRA NPS.^34^

Although there are obvious structural similarities to MEPIRAPIM, neither 5F-PY-PICA nor 5F-PY-PINACA had been described in the chemical literature prior to their detection on the NPS market. Pyrrolidine is a pendant structural feature of many psychoactive cathinones,^35^ but 1-(5-fluoropentyl)-1*H*-ind(az)ole-3-carboxamide is a common structural motif in SCRA NPS. The recent discovery in Spain of a phenethylamine featuring a popular SCRA functional group suggests that the design of NPS may not follow conventional rules for drug design, or that molecular hybridization is used in an ad hoc fashion to generate new compounds.^36^

Since nothing is known of the chemistry or pharmacology of 5F-PY-PICA and 5F-PY-PINACA, the aim of the present work was to prepare and analytically characterize these emergent SCRAs and several “prophetic” analogues, and to evaluate their cannabinoid receptor affinities and activities *in vitro* and *in vivo*. Related analogues arising from expansion (**7**, **8**), or scission (**9**, **10**) of the pyrrolidine ring were also prepared and evaluated to provide insight into structure-activity relationships (SARs) for this class. Additionally, several 5F-PY-PICA analogues featuring alternative 1-indole substituents commonly found in other SCRA classes were also prepared and characterized proactively. Specifically, analogues featuring a butyl (**11**), pentyl (**12**), 4-fluorobenzyl (**13**), and cyclohexylmethyl (**14**) subunit were synthesized, based on current SCRA design trends.

### Experimental. General chemical synthesis details

All reactions were performed under an atmosphere of nitrogen or argon unless otherwise specified. Dichloromethane, *N*,*N*-dimethylformamide (DMF), methanol, and toluene were anhydrous and used as purchased. Triethylamine was distilled over calcium hydride. All other commercially available reagents (Sigma-Aldrich) were used as purchased. Analytical thin layer chromatography (TLC) was performed using Merck aluminum-backed silica gel 60 F254 (0.2 mm) plates which were visualized using shortwave (254 nm) ultra-violet fluorescence. Flash chromatography was performed using Merck Kieselgel 60 (230–400 mesh) silica gel. Melting points were measured in open capillaries using an Laboratory Devices Mel-Temp II and are uncorrected. Nuclear magnetic resonance spectra were recorded at 298 K or 308 K using a Agilent 400 MHz spectrometer. The data are reported as chemical shift (*δ* ppm) relative to the residual protonated solvent resonance, relative integral, multiplicity (s = singlet, br = broad singlet, d = doublet, t = triplet, q = quartet, quin. = quintet, m = multiplet), coupling constants (*J* Hz) and assignment. Assignment of signals was assisted by COSY, DEPT, HSQC, and HMBC experiments where necessary. For compounds containing saturated heterocyclic amides (**5**–**8**, **11**–**14**), severe exchange broadening of the amide α- and β-carbon signals was observed, consistent with previous characterization of analogous systems.^37^

### General procedure for amidation of 1-alkyl-1*H*-indole-3-carboxylic acids

A solution of the appropriate carboxylic acid (0.5 mmol) in CH_2_Cl_2_ (1 mL) was treated with (COCl)_2_ (85 μL, 1.0 mmol, 2.0 equiv.) followed by DMF (1 drop). After stirring for 2 h, the solution was evaporated *in vacuo*, and the crude acid chloride was used immediately in the following step.

A cooled (0 °C) solution of the freshly prepared acid chloride (0.5 mmol) in CH_2_Cl_2_ (5 mL) was treated with Et_3_N (175 μL, 1.25 mmol, 2.5 equiv.), the appropriate amine (0.60 mmol, 1.2 equiv.), and stirred at ambient temperature for 14 h. The mixture was partitioned between CH_2_Cl_2_ (20 mL) and 1 M aq. HCl (10 mL). The layers were separated and the organic phase was washed with 1 M aq. HCl (2 × 10 mL), sat. aq. NaHCO_3_ (3 × 10 mL), brine (10 mL), dried (MgSO_4_), and the solvent evaporated under reduced pressure. The crude products were purified by flash chromatography unless stated otherwise.

### (1-(5-Fluoropentyl)-1*H*-indol-3-yl)(pyrrolidin-1-yl)methanone (5, 5F-PY-PICA)

Treating 1-(5-fluoropentyl)-1*H*-indole-3-carboxylic acid (125 mg, 0.5 mmol) with pyrrolidine (50 μL, 0.60 mmol, 1.2 equiv.) according to the general procedure afforded, following purification by flash chromatography (EtOAc), **5** as a white solid (135 mg, 89%). m.p. 66–68 °C; R_*f*_ 0.41 (EtOAc); ^1^H NMR (400 MHz, CDCl_3_): *δ* 8.15-8.12 (1H, m, CH), 7.41 (1H, s, CH), 7.34-.7.31 (1H, m, CH), 7.26 (1H, td, *J* = 7.5, 1.3 Hz, CH), 7.21 (1H, ddd, *J* = 7.9, 6.9, 1.2 Hz, CH), 4.41 (2H, dt, ^2^*J*_HF_ = 47.3 Hz, ^3^*J*_HH_ = 5.9 Hz, CH_2_F), 4.15 (2H, t, *J* = 7.1 Hz, CH_2_), 3.69 (4H, t, *J* = 6.6 Hz, CH_2_), 1.98-1.87 (6H, m, CH_2_), 1.77-1.64 (2H, m, CH_2_), 1.49-1.42 (2H, m, CH_2_); ^13^C NMR (100 MHz, CDCl_3_): *δ* 165.5 (CO), 135.9 (C_quat._), 129.5 (CH), 127.7 (C_quat._), 122.6 (CH), 122.5 (CH), 121.1 (CH), 111.7 (C_quat._), 109.5 (CH), 83.8 (d, ^1^*J*_CF_ = 165.2 Hz, CH_2_F), 47.7 (br, 2 × CH_2_), 46.7 (CH_2_), 30.1 (d, ^2^*J*_CF_ = 19.8 Hz, CH_2_), 29.8 (CH_2_), 25.5 (br, 2 × CH_2_), 23.0 (d, ^3^*J*_CF_ = 5.0 Hz, CH_2_); HRMS (ESI) *m/z* calcd for C18H23N2OF, 302.1794, found 302.1793. All physical properties and spectral data were consistent with those reported.^38^

### (1-(5-Fluoropentyl)-1*H*-indazol-3-yl)(pyrrolidin-1-yl)methanone (6, 5F-PY-PINACA)

To a solution of 1-(5-fluoropentyl)-1*H*-indazole-3-carboxylic acid (125 mg, 0.50 mmol) in DMF (2.5 mL) was added HOBt·H_2_O (84 mg, 0.55 mmol, 1.1 equiv.), EDC·HCl (125 mg, 0.65 mmol, 1.3 equiv.), pyrrolidine (42 μL, 0.50 mmol, 1.0 equiv.), and Et_3_N (140 μL, 1.00 mmol, 2.0 equiv.), and the mixture stirred at ambient temperature for 13 h. The mixture was poured onto H_2_O (200 mL), extracted with EtOAc (3 × 25 mL), and the combined organic were washed with H_2_O (2 × 25 mL), brine (25 mL), dried (MgSO_4_), and the solvent evaporated under reduced pressure. The crude material was purified using flash chromatography (hexane-EtOAc, 50:50) to give **6** as a colorless oil (137 mg, 90%). R_*f*_ 0.41 (hexane-EtOAc, 50:50); ^1^H NMR (400 MHz, CDCl_3_): *δ* 8.36 (1H, dt, *J* = 8.2, 1.0 Hz, CH), 7.40 (1H, t, *J* = 1.0 Hz, CH), 7.39 (1H, d, *J* = 1.0 Hz, CH), 7.26-7.22 (1H, m, CH), 4.409 (2H, t, *J* = 7.0 Hz, CH_2_), 4.408 (2H, dt, ^2^*J*_HF_ = 47.3 Hz, ^3^*J*_HH_ = 6.0 Hz, CH_2_F), 4.02 (2H, t, *J* = 6.6 Hz, CH_2_), 3.75 (2H, t, *J* = 6.7 Hz, CH_2_), 2.04-1.90 (6H, m, CH_2_), 1.79-1.66 (2H, m, CH_2_), 1.50-1.42 (2H, m, CH_2_); ^13^C NMR (100 MHz, CDCl_3_): *δ* 162.5 (CO), 140.1 (C_quat._), 139.2 (C_quat._), 126.7 (CH), 124.7 (C_quat._), 123.6 (CH), 122.2 (CH), 108.8 (CH), 83.8 (d, ^1^*J*_CF_ = 165.2 Hz, CH_2_F), 49.1 (CH_2_), 49.0 (CH_2_), 46.9 (CH_2_), 30.1 (d, ^2^*J*_CF_ = 19.8 Hz, CH_2_), 29.4 (CH_2_), 26.8 (CH_2_), 24.1 (CH_2_), 22.8 (d, ^3^*J*_CF_ = 5.2 Hz, CH_2_); HRMS (ESI) *m/z* calcd for C17H22N3OF, 303.1747, found 303.1746. All physical properties and spectral data were consistent with those reported.^39^

### (1-(5-Fluoropentyl)-1*H*-indol-3-yl)(piperidin-1-yl)methanone (7)

Treating 1-(5-fluoropentyl)-1*H*-indole-3-carboxylic acid (125 mg, 0.5 mmol) with piperidine (60 μL, 0.60 mmol, 1.2 equiv.) according to the general procedure gave, following purification by flash chromatography (hexane-EtOAc 50:50), **7** as a colorless oil (125 mg, 79%). R_*f*_ 0.56 (EtOAc); ^1^H NMR (400 MHz, CDCl_3_, 308 K): *δ* 7.71 (1H, ddd, *J* = 7.9, 1.4, 0.8 Hz, CH), 7.41 (1H, s, CH), 7.35-7.33 (1H, m, CH), 7.26-7.22 (1H, m, CH), 7.18 (1H, ddd, *J* = 8.0, 7.0, 1.1 Hz, CH), 4.41 (2H, dt, ^2^*J*_HF_ = 47.3 Hz, ^3^*J*_HH_ = 6.0 Hz, CH_2_F), 4.14 (2H, t, *J* = 7.1 Hz, CH_2_), 3.67-3.64 (4H, m, CH_2_), 1.92 (2H, quin., *J* = 7.5 Hz, CH_2_), 1.78-1.59 (8H, m, CH_2_), 1.51-1.43 (2H, m, CH_2_); ^13^C NMR (100 MHz, CDCl_3_, 308 K): *δ* 166.6 (CO), 135.9 (C_quat._), 130.1 (CH), 126.5 (C_quat._), 122.3 (CH), 121.0 (CH), 120.8 (CH), 111.4 (C_quat._), 109.8 (CH), 83.7 (d, ^1^*J*_CF_ = 165.3 Hz, CH_2_F), 46.6 (CH_2_), 46.3 (br, 2 × CH_2_), 30.1 (d, ^2^*J*_CF_ = 19.8 Hz, CH_2_), 29.8 (CH_2_), 26.5 (br, 2 × CH_2_), 24.9 (CH_2_), 23.0 (d, ^3^*J*_CF_ = 5.0 Hz, CH_2_); HRMS (ESI) *m/z* calcd for C19H25N2OF, 316.1951, found 316.1953.

### Azepan-1-yl(1-(5-fluoropentyl)-1*H*-indol-3-yl)methanone (8)

Treating 1-(5-fluoropentyl)-1*H*-indole-3-carboxylic acid (125 mg, 0.5 mmol) with homopiperidine (65 μL, 0.60 mmol, 1.2 equiv.) according to the general procedure gave, following purification by flash chromatography (hexane-EtOAc 50:50), **8** (130 mg, 79%) as a colorless oil. R_*f*_ 0.63 (EtOAc); ^1^H NMR (400 MHz, CDCl_3_, 308 K): *δ* 7.82 (1H, dt, *J* = 7.8, 1.1 Hz, CH), 7.34-7.31 (2H, m, CH, overlapping), 7.24 (1H, ddd, *J* = 8.2, 7.0, 1.3 Hz, CH), 7.17 (1H, ddd, *J* = 8.0, 7.0, 1.1 Hz, CH), 4.40 (2H, dt, ^2^*J*_HF_ = 47.3 Hz, ^3^*J*_HH_ = 6.0 Hz, CH_2_F), 4.13 (2H, t, *J* = 7.1 Hz, CH_2_), 3.70-3.67 (4H, m, CH_2_), 1.90 (2H, quin., *J* = 7.5 Hz, CH_2_), 1.83-1.58 (10H, m, CH_2_), 1.49-1.42 (2H, m, CH_2_); ^13^C NMR (100 MHz, CDCl_3_, 308 K): *δ* 167.3 (CO), 135.8 (C_quat._), 128.5 (CH), 127.2 (C_quat._), 122.4 (CH), 121.5 (CH), 120.7 (CH), 111.5 (C_quat._), 109.6 (CH), 83.7 (d, ^1^*J*_CF_ = 165.3 Hz, CH_2_F), 48.1 (br, 2 × CH_2_), 46.6 (CH_2_), 30.0 (d, ^2^*J*_CF_ = 19.8 Hz, CH_2_), 29.8 (CH_2_), 28.7 (br, 2 × CH_2_), 27.6 (br, 2 × CH_2_), 23.0 (d, ^3^*J*_CF_ = 5.0 Hz, CH_2_); HRMS (ESI) *m/z* calcd for C20H27N2OF, 330.2107, found 330.2113.

### 1-(5-Fluoropentyl)-*N*,*N*-diethyl-1*H*-indole-3-carboxamide (9)

Treating 1-(5-fluoropentyl)-1*H*-indole-3-carboxylic acid (125 mg, 0.5 mmol) with diethylamine (60 μL, 0.60 mmol, 1.2 equiv.) according to the general procedure gave, following purification by flash chromatography (EtOAc), **9** (125 mg, 82%) as a colorless oil. R_*f*_ 0.61 (EtOAc); ^1^H NMR (400 MHz, CDCl_3_, 308 K): *δ* 7.77 (1H, ddd, *J* = 7.9, 1.3, 0.8 Hz, CH), 7.33 (1H, dt, *J* = 8.2, 0.9 Hz, CH, overlapping), 7.32 (1H, s, CH, overlapping), 7.24 (1H, ddd, *J* = 8.2, 7.0, 1.3 Hz, CH), 7.18 (1H, ddd, *J* = 7.9, 7.0, 1.0 Hz, CH), 4.41 (2H, dt, ^2^*J*_HF_ = 47.3, ^3^*J*_HH_ = 5.9 Hz, CH_2_F), 4.15 (2H, t, *J* = 7.1 Hz, NCH_2_), 3.57 (4H, q, *J* = 7.1 Hz, 2 × NCH_2_), 1.91 (2H, quin., *J* = 7.5 Hz, CH_2_), 1.78-1.65 (2H, m, CH_2_), 1.50-1.43 (2H, m, CH_2_), 1.22 (6H, t, *J* = 7.1 Hz, 2 × CH_3_); ^13^C NMR (100 MHz, CDCl_3_, 308 K): *δ* 167.0 (CO), 135.9 (C_quat._), 128.3 (CH), 127.1 (C_quat._), 122.5 (CH), 121.2 (CH), 120.7 (CH), 111.6 (C_quat._), 109.7 (CH), 83.8 (d, ^1^*J*_CF_ = 165.2 Hz, CH_2_F), 46.6 (CH_2_), 41.5 (br, 2 × CH_2_), 30.1 (d, ^2^*J*_CF_ = 19.8 Hz, CH_2_), 29.9 (CH_2_), 23.0 (d, ^3^*J*_CF_ = 5.0 Hz, CH_2_), 14.0 (2 × CH_3_); HRMS (ESI) *m/z* calcd for C18H25N2OF, 304.1951, found 304.1953.

### 1-(5-Fluoropentyl)-*N*,*N*-dimethyl-1*H*-indole-3-carboxamide (10)

Treating 1-(5-fluoropentyl)-1*H*-indole-3-carboxylic acid (125 mg, 0.5 mmol) with a solution of dimethylamine in THF (2 M, 300 μL, 0.60 mmol, 1.2 equiv.) according to the general procedure gave, following purification by flash chromatography (EtOAc), **10** (116 mg, 84%) as a colorless oil. R_*f*_ 0.31 (EtOAc); ^1^H NMR (400 MHz, CDCl_3_, 308 K): *δ* 7.79 (1H, ddd, *J* = 7.9, 1.4, 0.9 Hz, CH), 7.41 (1H, s, CH), 7.34 (1H, dt, *J* = 8.2, 0.9 Hz, CH), 7.25 (1H, ddd, *J* = 8.2, 6.9, 1.3 Hz, CH), 7.20 (1H, ddd, *J* = 7.9, 6.9, 1.3 Hz, CH), 4.42 (2H, dt, *J* = 47.3, 5.9 Hz, CH_2_F), 4.15 (2H, t, *J* = 7.1 Hz, NCH_2_), 3.16 (6H, s, 2 × CH_3_), 1.95 (2H, quin. *J* = 7.5 Hz, CH_2_), 1.78-1.65 (2H, m, CH_2_), 1.51-1.43 (2H, m, CH_2_); ^13^C NMR (100 MHz, CDCl_3_, 308 K): *δ* 167.8 (CO), 135.9 (C_quat._), 130.0 (CH), 126.9 (C_quat._), 122.5 (CH), 121.6 (CH), 120.9 (CH), 111.1 (C_quat._), 109.8 (CH), 83.8 (d, ^1^*J*_CF_ = 165.2 Hz, CH_2_F), 46.7 (NCH_2_), 37.8 (br, 2 × CH_3_), 30.1 (d, ^2^*J*_CF_ = 19.8 Hz, CH_2_), 29.9 (CH_2_), 23.0 (d, ^3^*J*_CF_ = 5.0 Hz, CH_2_); HRMS (ESI) *m/z* calcd for C16H21N2OF, 276.1638, found 276.1643.

### (1-Butyl-1*H*-indol-3-yl)(pyrrolidin-1-yl)methanone (11)

Treating 1-butyl-1*H*-indole-3-carboxylic acid (109 mg, 0.5 mmol) with pyrrolidine (45 μL, 0.55 mmol, 1.1 equiv.) according to the general procedure gave, following purification by flash chromatography (EtOAc), **11** (113 mg, 84%) as colorless crystals. m.p. 102.5–104.5 °C; R_*f*_ 0.41 (EtOAc); ^1^H NMR (400 MHz, CDCl_3_, 308 K): *δ* 8.14 (1H, ddd, *J* = 7.8, 1.5, 0.8 Hz, CH), 7.41 (1H, s, CH), 7.34-7.31 (1H, m, CH), 7.24 (1H, ddd, *J* = 8.1, 6.9, 1.4 Hz, CH), 7.20 (1H, ddd, *J* = 8.1, 6.8, 1.4 Hz, CH), 4.12 (2H, t, *J* = 7.1 Hz, CH_2_), 3.70-3.67 (4H, m, CH_2_), 1.97-1.90 (4H, m, CH_2_), 1.86-1.79 (2H, m, CH_2_), 1.40-1.31 (2H, m, CH_2_), 0.94 (3H, t, *J* = 7.4 Hz, CH_3_); ^13^C NMR (100 MHz, CDCl_3_, 308 K): *δ* 165.5 (CO), 135.9 (C_quat._), 129.6 (CH), 127.7 (C_quat._), 122.5 (CH), 122.4 (CH), 120.9 (CH), 111.4 (C_quat._), 109.5 (CH), 47.8 (br, 2 × CH_2_), 46.6 (CH_2_), 32.2 (CH_2_), 25.7 (br, 2 × CH_2_), 20.2 (CH_2_), 13.7 (CH_3_); HRMS (ESI) *m/z* calcd for C17H22N2O, 270.1732, found 270.1734.

### (1-Pentyl-1*H*-indol-3-yl)(pyrrolidin-1-yl)methanone (12)

Treating 1-pentyl-1*H*-indole-3-carboxylic acid (116 mg, 0.5 mmol) with pyrrolidine (45 μL, 0.55 mmol, 1.1 equiv.) according to the general procedure gave, following purification by flash chromatography (EtOAc), **12** (89 mg, 63%) as colorless crystals. m.p. 57–59 °C; R_*f*_ 0.37 (EtOAc); ^1^H NMR (400 MHz, CDCl_3_, 308 K): *δ* 8.15-8.12 (1H, m, CH), 7.41 (1H, s, CH), 7.34-7.32 (1H, m, CH), 7.25 (1H, ddd, *J* = 8.1, 7.0, 1.4 Hz, CH), 7.20 (1H, ddd, *J* = 8.0, 6.9, 1.3 Hz, CH), 4.12 (2H, t, *J* = 7.2 Hz, CH_2_), 3.75-3.64 (4H, m, CH_2_), 1.99-1.90 (4H, m, CH_2_), 1.86 (2H, quin., *J* = 7.2 Hz, CH_2_), 1.39-1.27 (4H, m, CH_2_), 0.89 (3H, t, *J* = 7.0 Hz, CH_3_); ^13^C NMR (100 MHz, CDCl_3_, 308 K): *δ* 165.6 (CO), 135.9 (C_quat._), 129.6 (CH), 127.7 (C_quat._), 122.5 (CH), 122.4 (CH), 121.0 (CH), 111.5 (C_quat._), 109.6 (CH), 47.9 (br, 2 × CH_2_), 46.9 (CH_2_), 29.9 (CH_2_), 29.2 (CH_2_), 25.6 (br, 2 × CH_2_), 22.4 (CH_2_), 14.0 (CH_3_); HRMS (ESI) *m/z* calcd for C18H24N2O, 284.1889, found 284.1889.

### (1-(4-Fluorobenzyl)-1*H*-indol-3-yl)(pyrrolidin-1-yl)methanone (13)

Treating 1-(4-fluorobenzyl)-1*H*-indole-3-carboxylic acid (135 mg, 0.5 mmol) with pyrrolidine (45 μL, 0.55 mmol, 1.1 equiv.) according to the general procedure gave, following purification by flash chromatography (EtOAc), **13** (129 mg, 80%) as a white solid. m.p. 159–161 °C; R_*f*_ **0.33 (EtOAc); 1H NMR (400 MHz, CDCl_3_, 308 K):***δ* **8.17-8.13 (1H, m, CH), 7.41 (1H, m, CH), 7‥27-7.20 (3H, m, CH), 7.12-7.07 (2H, m, CH), 7.02-6.96 (2H, m, CH), 5.30 (2H, s, CH_2_), 3.67 (4H, br s, CH_2_), 1.97-1.90 (4H, m, CH_2_); ^13^C NMR (100 MHz, CDCl_3_, 308 K):***δ* 165.3 (CO), 162.6 (d, ^1^*J*_CF_ = 247.0 Hz, CF), 136.1 (C_quat._), 132.4 (d, ^4^*J*_CF_ = 3.5 Hz, C_quat._), 129.7 (CH), 128.7 (d, ^3^*J*_CF_ = 8.1 Hz, CH), 127.8 (C_quat._), 123.0 (CH), 122.5 (CH), 121.4 (CH), 116.0 (d, ^2^*J*_CF_ = 21.9 Hz, CH), 112.4 (C_quat._), 109.8 (CH), 49.9 (CH_2_), 48.0 (br, 2 × CH_2_), 25.7 (br, 2 × CH_2_); HRMS (ESI) *m/z* calcd for C20H19N2OF, 322.1481, found 322.1484.

### (1-(Cyclohexylmethyl)-1*H*-indol-3-yl)(pyrrolidin-1-yl)methanone (14)

Treating 1-(cyclohexylmethyl)-1*H*-indole-3-carboxylic acid (129 mg, 0.5 mmol) with pyrrolidine (45 μL, 0.55 mmol, 1.1 equiv.) according to the general procedure gave, following purification by flash chromatography (EtOAc), **14** (130 mg, 84%) as a white solid. m.p. 156–158 °C; R_*f*_ 0.39 (EtOAc); 1H NMR (400 MHz, CDCl_3_, 308 K): *δ* 8.14 (1H, ddd, *J* = 7.8, 1.5, 0.8 Hz, CH), 7.37 (1H, s, CH), 7.34-7.31 (1H, m, CH), 7.27-7.22 (1H, m, CH), 7.21-7.18 (1H, m, CH), 3.95 (2H, d, *J* = 7.2 Hz, CH_2_), 3.73-3.66 (4H, m, CH_2_), 1.97-1.93 (4H, m, CH_2_), 1.90-1.81 (1H, m, CH), 1.76-1.61 (5H, m, CH_2_), 1.25-1.10 (3H, m, CH_2_), 1.06-0.95 (2H, m, CH_2_); ^13^C NMR (100 MHz, CDCl_3_, 308 K): *δ* 165.6 (CO), 136.3 (C_quat._), 130.4 (CH), 127.7 (C_quat._), 122.5 (CH), 122.4 (CH), 121.0 (CH), 111.2 (C_quat._), 109.8 (CH), 53.4 (CH_2_), 47.8 (br, 2 × CH_2_), 38.8 (CH), 31.2 (2 × CH_2_) 26.4 (CH_2_), 25.9 (br, 2 × CH_2_), 25.8 (2 × CH_2_); HRMS (ESI) *m/z* calcd for C20H26N2O, 310.2045, found 310.2048.

### General procedure for synthesis of 1-alkyl-1*H*-indole-3-carboxylic acids

To a cooled (0 °C) suspension of sodium hydride (60% dispersion in mineral oil, 120 mg, 3.00 mmol, 2.0 equiv.) in DMF (1.5 mL) was added dropwise a solution of indole (**15**, 176 mg, 1.50 mmol) in DMF (0.2 mL) and the mixture allowed to stir at ambient temperature for 10 min. The mixture was cooled (0 °C), treated dropwise with the appropriate alkyl bromide (1.58 mmol, 1.05 equiv.), and stirred at ambient temperature for 1 h. The mixture was cooled (0 °C), treated portionwise with trifluoroacetic anhydride (0.521 mL, 3.75 mmol, 2.5 equiv.), and stirred at ambient temperature for 1 h. The solution was poured portionwise onto vigorously stirred ice-water (90 mL) until precipitation was complete, and the red-pink solid was filtered and air dried overnight.

To a refluxing solution of potassium hydroxide (278 mg, 4.95 mmol, 3.3 equiv.) in methanol (0.55 mL) was added portionwise a solution of the crude 1-alkyl-3-trifluoroacetyl-1*H*-indole in toluene (1.25 mL) and the solution heated at reflux for 2 h. The solution was cooled to ambient temperature and partitioned between 1 M aq. NaOH (40 mL) and Et_2_O (5 mL). The layers were separated and the aqueous layer was adjusted to pH 1 with 10 M aq. HCl. The aqueous phase was extracted with Et_2_O (3 × 10 mL), dried (Na_2_SO_4_), and solvent evaporated under reduced pressure. The crude products were recrystallized from *i*-PrOH unless otherwise stated.

### 1-Butyl-1*H*-indole-3-carboxylic acid (16)

Subjecting 1-bromobutane (170 μL, 1.58 mmol) to the general procedure above gave **16** as a colorless crystalline solid (201 mg, 62% over 2 steps). m.p. 134–135 °C; ^1^H NMR (400 MHz, CDCl_3_, 298 K): *δ* 8.28-8.23 (1H, m, CH), 7.93 (1H, s, CH), 7.42-7.37 (1H, m, CH), 7.33-7.29 (2H, m, CH), 4.17 (2H, d, *J* = 7.1 Hz, CH_2_), 1.92-1.84 (2H, m, CH_2_), 1.43-1.33 (2H, m, CH_2_), 0.97 (3H, t, *J* = 7.4 Hz, CH_3_); All spectral data were consistent with those previously reported.^40^

### 1-Pentyl-1*H*-indole-3-carboxylic acid (17)

Subjecting 1-bromopentane (195 μL, 1.58 mmol) to the general procedure gave **17** as a colorless crystalline solid (192 mg, 55% over 2 steps). m.p. 106–108 °C (lit. m.p. 106–108 °C); ^1^H NMR (400 MHz, CDCl_3_, 298 K): *δ* 8.30-8.26 (1H, m, CH), 7.94 (1H, s, CH), 7.41-7.38 (1H, m, CH), 7.34-7.29 (2H, m, CH), 4.16 (2H, d, *J* = 7.2 Hz, CH_2_), 1.90 (2H, quin., *J* = 7.2 Hz, CH_2_), 1.40-1.30 (4H, m, CH_2_), 0.91 (3H, t, *J* = 7.4 Hz, CH_3_). All physical properties and spectral data were consistent with those reported.^41^

### 1-(5-Fluoropentyl)-1*H*-indole-3-carboxylic acid (18)

Subjecting 1-bromo-5-fluoropentane (195 μL, 1.58 mmol) to the general procedure gave **18** as a colorless crystalline solid (247 mg, 66% over 2 steps). m.p. 120–122 °C (lit. m.p. 120–122 °C); ^1^H NMR (400 MHz, CDCl_3_, 298 K): *δ* 8.29-8.25 (1H, m, CH), 7.93 (1H, s, CH), 7.41-7.37 (1H, m, CH), 7.34-7.30 (2H, m, CH), 4.43 (2H, dt, ^2^*J*_CF_ = 47.3 Hz, ^3^*J*_HH_ = 5.9 Hz, CH_2_F), 4.18 (2H, d, *J* = 7.1 Hz, CH_2_), 1.98-1.91 (2H, m, CH_2_), 1.79-1.66 (2H, m, CH_2_), 1.52-1.44 (2H, m, CH_2_). All physical properties and spectral data were consistent with those reported.^42^

### 1-(4-Fluorobenzyl)-1*H*-indole-3-carboxylic acid (19)

Subjecting 4-fluorobenzyl bromide (195 μL, 1.58 mmol) to the general procedure gave **19** as a colorless crystalline solid (256 mg, 63% over 2 steps). m.p. 208–210 °C (lit. m.p. 207–209 °C); ^1^H NMR (400 MHz, CDCl_3_, 298 K): *δ* 8.26-8.24 (1H, m, CH), 7.92 (1H, s, CH), 7.34-7.26 (3H, m, CH), 7.18-7.14 (2H, m, CH), 7.06-7.00 (2H, m, CH), 5.33 (2H, s, CH_2_). All physical properties and spectral data were consistent with those reported.^43^

### 1-(Cyclohexylmethyl)-1*H*-indole-3-carboxylic acid (20)

Subjecting (bromomethyl)cyclohexane (220 μL, 1.58 mmol) to the general procedure gave **20** as a colorless crystalline solid (174 mg, 45%). m.p. 180–182 °C (lit. m.p. 180–182 °C); 1H NMR (400 MHz, CDCl_3_, 298 K): *δ* 8.27-8.23 (1H, m, CH), 7.89 (1H, s, CH), 7.40-7.36 (1H, m, CH), 7.32-7.28 (2H, m, CH), 3.99 (2H, d, *J* = 7.2 Hz, CH_2_), 1.95-1.85 (1H, m, CH), 1.76-1.62 (5H, m, CH_2_), 1.26-1.15 (3H, m, CH_2_), 1.06-0.97 (2H, m, CH_2_). All physical properties and spectral data were consistent with those reported.^43^

### Methyl 1-(5-fluoropentyl)-1*H*-indazole-3-carboxylate (22)

To a cooled (0 °C) solution of methyl 1*H*-indazole-3-caboxylate (**21**, 176 mg, 1.00 mmol, 1.0 equiv.) in THF (5.0 mL) was added dropwise a 1 M solution of potassium tert-butoxide in THF (1.1 mL, 1.1 mmol, 1.1 equiv.), and the solution stirred at ambient temperature for 1 h. The solution was cooled (0 °C), treated dropwise with 1-bromo-5-fluoropentane (130 μL, 1.05 mmol, 1.05 equiv.), and stirred at ambient temperature for 14 h. The mixture was partitioned between EtOAc (10 mL) and H_2_O (30 mL) and the layers separated. The aqueous layer was extracted with EtOAc (3 × 10 mL), and the combined organic phases were washed with brine (10 mL), dried (Na_2_SO_4_), and the solvent evaporated under reduced pressure affording, following purification by flash chromatography (hexane-EtOAc, 90:10), **22** (214 mg, 81%) as a colorless oil. R_*f*_ 0.59 (hexane-EtOAc, 80:20); ^1^H NMR (400 MHz, CDCl_3_, 298 K): *δ* 8.20 (1H, dt, J = 8.2, 1.0 Hz), 7.45-7.38 (2H, m), 7.28 (1H, ddd, *J* = 8.2, 6.4, 1.3 Hz), 4.45 (2H, t, *J* = 7.2 Hz, CH_2_), 4.37 (2H, dt, ^2^*J*_HF_ = 47.3 Hz, ^3^*J*_HH_ = 6.0 Hz, CH_2_F), 4.00 (3H, s, CH_3_), 2.02-1.95 (2H, m, CH_2_), 1.74-1.61 (2H, m, CH_2_), 1.46-1.38 (2H, m, CH_2_). All physical properties and spectral data were consistent with those reported.^13^

### 1-(5-fluoropentyl)-1*H*-indazole-3-carboxylic acid (23)

To a solution of **22** (198 mg, 0.75 mmol) in MeOH (7.5 mL) was added 1 M aq. NaOH (3.0 mL) in a single portion, and the solution stirred at ambient temperature for 16 h. The solvent was evaporated under reduced pressure, the crude material dissolved in H_2_O (30 mL), and the pH adjusted to 2 with 10 M aq. HCl. The aqueous phase was extracted with EtOAc (3 × 10 mL), and the combined organic layers were washed with brine (5 mL), dried (Na_2_SO_4_), and the solvent evaporated under reduced pressure to afford **23** (182 mg, 97%). m.p. 80-82 °C; (lit. m.p. 80-82 °C); ^1^H NMR (400 MHz, CDCl_3_, 298 K): *δ* 8.27 (1H, dt, *J* = 8.2, 1.0 Hz, CH), 7.52-7.46 (2H, m, CH), 7.37 (1H, ddd, *J* = 8.2, 6.2, 1.7 Hz, CH), 4.53 (2H, t, J = 7.2 Hz, CH2), 4.44 (2H, dt, ^2^*J*_HF_ = 47.3 Hz, ^3^*J*_HH_ = 5.9 Hz, CH_2_F), 2.10-2.02 (2H, m, CH_2_), 1.80-1.67 (2H, m, CH_2_), 1.52-1.44 (2H, m, CH_2_). All physical properties and spectral data were consistent with those reported.^13^

### QTOF LC-MS/MS acquisition and analysis method

All samples were analyzed using an Agilent LC 1260 Infinity Binary Liquid Chromatograph (LC) System attached to an Agilent 6550 iFunnel Quadrupole Time-of-Flight Mass Spectrometer (QTOF/MS) 6550 (Agilent Technologies, Santa Clara, CA). An Agilent jet stream electrospray ionization (ESI) source with a dual nebulizer that allows constant introduction of reference mass during sample run was used to ionize sample organic components in positive mode.

Each sample was prepared from a crystalline or resin aliquot of purified synthetic product and diluted to a final concentration of 100 and 500 ng/mL in 10% LC-MS grade acetonitrile (Honeywell B&J, Muskegon, MI); two concentrations were injected to elucidate potential solvent impurities or instrument artifacts. In each sample run, 2.5 μL was injected into an Agilent Poroshell 120 C-18 column (2.1 × 100 mm, 2.7 μm) maintained at 50 °C. Chromatographic separation was achieved by gradient elution using LC-MS grade water (Honeywell B&J, Muskegon, MI) with 0.05% formic acid and 5 mM ammonium formate as mobile phase A, and acetonitrile with 0.05% formic acid as mobile phase B. The elution gradient used was 0–0.5 min = 5% B; 1.5 min = 30% B; 4.5 min = 70% B; 7.5 min = 100% B; 7.5–10 min = 100% B; and 10.01–12 min = 5% B.

Ionization of chromatographic eluates was induced on the QTOF/MS using an ESI source in the positive mode operated under the following conditions: gas temperature at 225 °C; sheath gas temperature at 350 °C; drying gas flow at 14 L/min; sheath gas flow at 11 L/min; nebulizer pressure at 40 psig; voltage cap at 3000 V; and nozzle voltage at 500 V. Data acquisition was run at 2 GHz in extended dynamic range mode. Both TOF/MS and MS/MS spectra were collected in automated MS/MS mode using 500 arbitrary units as threshold for inducing MS/MS data collection. An active exclusion was used after 1 spectra, with a release time of 0.05 mins. Each sample was run in triplicate.

The total ion chromatogram (TIC) obtained from the LC-QTOF/MS run was analyzed using Agilent MassHunter Qualitative Analysis software (Agilent Technologies, Sta. Clara, CA). A search was done using the chemical formula of the expected material to confirm the identity and measure retention time of the major chromatogram peak. To confirm, the following criteria were imposed for a compound match: mass error ≤ 10 ppm; target score ≥ 70 (indication of isotopic pattern match) for peaks that did not exhibit detector saturation; and the presence of at least one expected fragment ion peak in its MS/MS spectra. MS/MS spectra were captured to assess the unique fragmentation of each parent cannabinoid.

### *In vitro* cannabinoid receptor binding experiments

Experiments utilized human CB_1_ or CB_2_ tagged at the *N*-terminus with three haemagglutinin sequences (HA-hCB_1_, HA-hCB_2_) stably transfected into HEK 293.^44,45^ HEK 293 were cultivated in Dulbecco’s Modified Eagle’s Medium (DMEM) supplemented with 10% fetal bovine serum (FBS) and zeocin, (250 μg/mL). Cells were maintained in 5% CO_2_ at 37 °C in a humidified atmosphere. Cells were grown in 75 cm^2^ flasks and passaged when 80-90% confluent.

HEK 293-hCB_1_ or HEK293-hCB_2_ cells were grown to 90–100% confluence in 175 cm^2^ flasks and harvested in ice-cold phosphate buffered saline (PBS) with 5 mM EDTA. Cells were centrifuged at 200 × g for 10 minutes and the pellet frozen at −80 °C until required. Pellets were thawed with Tris-sucrose buffer (50 mM Tris-HCl, pH 7.4, 200 mM sucrose, 5 mM MgCl_2_, 2.5 mM EDTA) and homogenized with a glass homogenizer. The homogenate was centrifuged at 1000 × g for 10 minutes at 4 °C and the pellet discarded. The supernatant was then centrifuged at 27 000 × g for 30 minutes at 4 °C. The final pellet was resuspended in a minimal volume of Tris-sucrose buffer, and aliquoted then stored at −80 °C. Protein concentration was determined using the DC protein assay kit (Bio-Rad, Hercules, CA, USA) following the manufacturers protocol. Initial compound screening involved resuspension of cell membranes (5-10 μg/point) in binding buffer (50 mM HEPES, 1 mM MgCl2, 1 mM CaCl2, 0.2% (w/v) bovine serum albumin (BSA, ICP Bio, New Zealand, pH 7.4) and incubation with [3H]SR141716A (1.25 nM, specific activity 55 Ci/mmol) for CB_1_ or [^3^H]CP55,940 (1.0 nM, specific activity 175 Ci/mmol) for CB_2_ (PerkinElmer, Waltham, MA, USA) and 10 μM of the test compound at 30 °C for 60 minutes. Test compounds were all dissolved in DMSO at 10 mM, and DMSO addition was controlled for in all assay points. GF/C Harvest Plates (PerkinElmer) were pre-soaked for one hour in 0.1% polyethylenimine and then washed with 200 μL ice cold wash buffer (50 mM HEPES pH 7.4 500 mM NaCl, 0.1 % BSA) prior to filtration of samples, which were then subject to 3 additional 200 μL washes in ice cold wash buffer. Harvest plates were dried overnight at 24 °C, and then 50 μL of scintillation fluid was added to each well and plates were read 30 minutes later for 2 minutes per well in a Microbeta Trilux (PerkinElmer). Screening assays were performed in duplicate, repeating assays for each mediator 3 or more times. Non-specific binding was defined as binding that occurred in the presence of 10 μM CP 55,940. Compounds that produced >60% displacement at 10 μM were further characterized to define affinity. Competitive binding curves were produced by incubating the membranes and radioligand with a range of concentrations of test compound. Data was analyzed using GraphPad Prism, curves are generated using a One site-fit *K*_i_ Competitive Binding function with *K*_d_ of [^3^H]SR141716A constrained to 1 nM at CB_1_, and [^3^H]CP 55,940 to 1 nM at CB_2_.

### *In vitro* cannabinoid receptor functional activity assay

Mouse AtT20 neuroblastoma cells stably transfected with human CB_1_ or human CB_2_ have been previously described.^43^ and were cultured in Dulbecco’s modified Eagle’s medium (DMEM) containing 10% fetal bovine serum (FBS), 100 U penicillin/streptomycin, and 80 μg/mL hygromycin (InvivoGen, San Diego, CA, USA). Cells were passaged at 80% confluency as required. Cells for assays were grown in 75 cm2 flasks and used at 90% confluence. The day before the assay cells were detached from the flask with trypsin/EDTA (Sigma) and resuspended in 10 mL of Leibovitz’s L-15 media supplemented with 1% FBS, 100 U penicillin/streptomycin and 15 mM glucose. The cells were plated in volume of 90 μL in black walled, clear bottomed 96-well microplates (Corning) which had been precoated with poly-L-lysine (Sigma, Australia). Cells were incubated overnight at 37 °C in ambient CO_2_.

Membrane potential was measured using a FLIPR Membrane Potential Assay kit (blue) from Molecular Devices, as described previously.^46^ The dye was reconstituted with assay buffer of composition (mM): NaCl 145, HEPES 22, Na_2_HPO_4_ 0.338, NaHCO_3_ 4.17, KH_2_PO_4_ 0.441, MgSO_4_ 0.407, MgCl_2_ 0.493, CaCl_2_ 1.26, glucose 5.56 (pH 7.4, osmolarity 315 ± 5). Prior to the assay, cells were loaded with 90 μL/well of the dye solution without removal of the L-15, giving an initial assay volume of 180 μL/well. Plates were then incubated at 37 °C at ambient CO_2_ for 45 min. Fluorescence was measured using a FlexStation 3 (Molecular Devices) microplate reader with cells excited at a wavelength of 530 nm and emission measured at 565 nm. Baseline readings were taken every 2 s for at least 2 min, at which time either drug or vehicle was added in a volume of 20 μL. The background fluorescence of cells without dye or dye without cells was negligible. Changes in fluorescence were expressed as a percentage of baseline fluorescence after subtraction of the changes produced by vehicle addition. The final concentration of DMSO was 0.1%.

Data were analyzed with PRISM (GraphPad Software Inc., San Diego, CA), using four-parameter nonlinear regression to fit concentration-response curves. In all plates, a maximally effective concentration of CP 55,940 was added to allow for normalization between assays.

### *In vivo* pharmacological assessment of 5F-PY-PICA and 5F-PY-PINACA

Two cohorts of 4 adult male C57BL/6J mice (Animal Resources Centre, Perth, Australia) were used for biotelemetric assessment of body temperature. The mice weighed between 20.5 and 28.5 g on arrival, and were singly housed in a climate-controlled testing room (23 ± 1 °C) on a 12 h light/dark cycle (lights on from 07:00 to 19:00). Water and standard rodent chow were provided ad libitum. All experiments were approved by The University of Sydney Animal Ethics Committee.

Biotelemetry transmitters (model TA-F10, Data Sciences International, St. Paul, MN) were implanted as previously described.^13^ The transmitter was implanted according to the manufacturers protocol into the peritoneal cavity following anesthetization with isoflurane (3% induction, 1-2% maintenance). The wound was sutured closed and data collection commenced after 10 days of recovery.

The mice were habituated over multiple days to injections of vehicle (a solution composed of 7.8% polysorbate 80 and 92.2% physiological saline). Injection always occurred at a set time of day (10:00 am). Each cohort received injections of each compound in an ascending dose sequence (0.3, 1, 3, and 10 mg/kg). This ascending sequence was used in order to minimize the risk posed to the animals in assessing hitherto untested compounds. Two washout (drug free) days were given between each dose to limit development of tolerance or carryover effects.

Body temperature data was gathered continuously at 1000 Hz and organized into 15 minute bins using Dataquest A.R.T. software (version 4.3, Data Sciences International, St. Paul, MN), and analysed using PRISM (version 7, Graphpad Software Inc., San Diego, CA).^13^

## Results and discussion

The synthesis of 5F-PY-PICA (**5**) and related analogues **7**–**15** is shown in Figure 2. Using a general procedure, indole (**15**) was first alkylated with the appropriate alkyl bromide, converted to the corresponding 3-trifluoroacetylindole, and hydrolyzed to the carboxylic acid (**16–20**). The relevant carboxylic acid **16–20** was converted to the corresponding acid chloride using oxalyl chloride, and treated with pyrrolidine, piperidine, azepane, diethylamine, or dimethylamine to furnish amides **5** and **7**–**14**.

**Figure 2.**
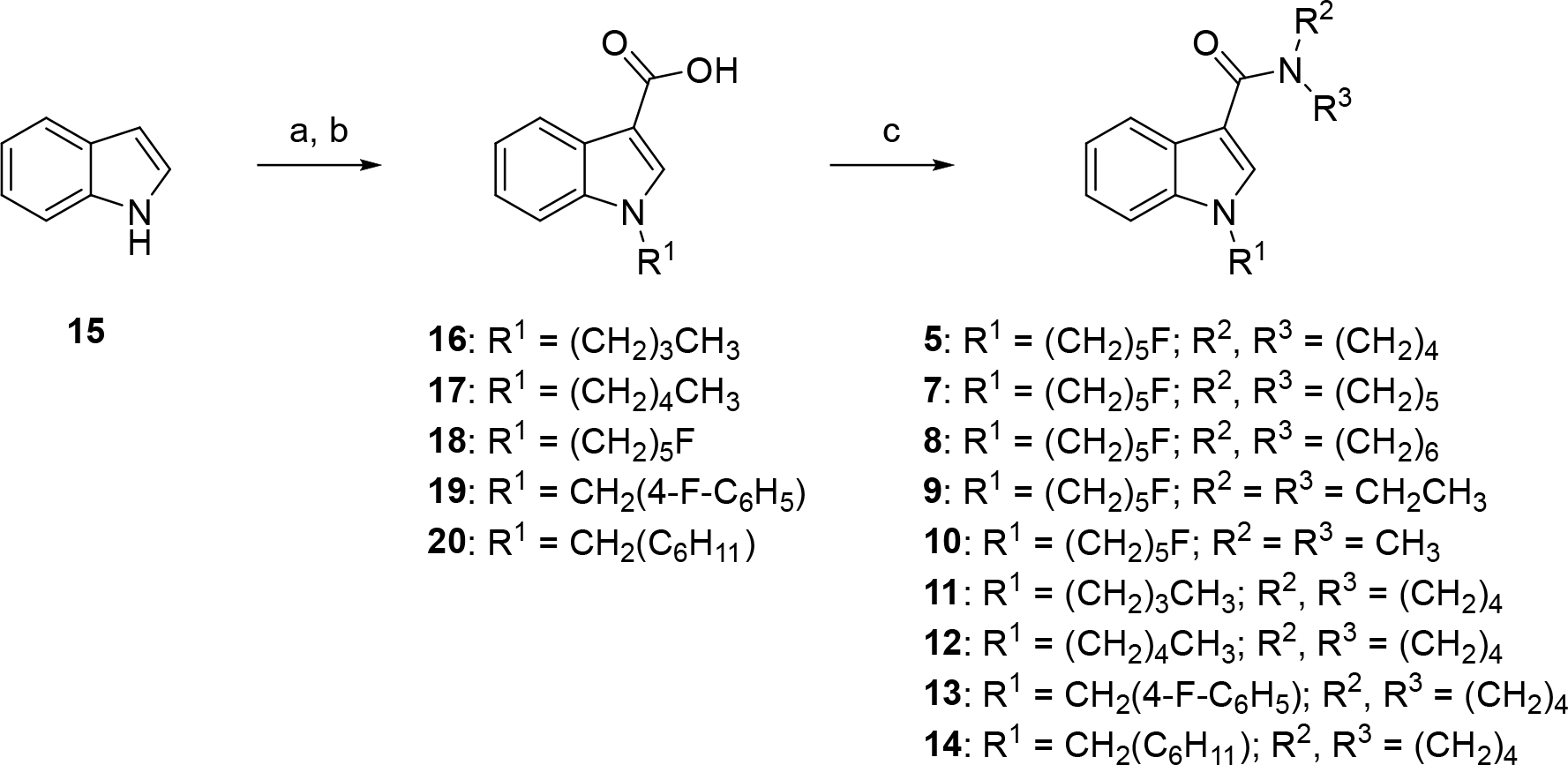
Reagents and conditions: (a)(i) NaH, R^1^Br, DMF, 0 °C-rt, 1 h; (ii) (CF_3_CO)_2_O, 0 °C-rt, 1 h; (b) KOH, MeOH, PhMe, reflux, 2 h, 45-66% over 3 steps; (c) (COCl)_2_, DMF (cat.), CH_2_Cl_2_, 0 °C-rt, 2 h; (d) R^2^R^3^NH, Et_3_N, CH_2_Cl_2_, 0 °C–rt, 14 h, 63-89%.

The synthesis of 5F-PY-PINACA (**6**) required a slightly different route and is depicted in Figure 3. Methyl 1*H*-indazole-3-carboxylate (**21**) was deprotonated with potassium *tert*-butoxide, and then treated with 1-bromo-5-fluoropentane, to regioselectively yield the desired 1-alkylated indole-3-carboxylate (**22**) as previously described.^13^ Saponification of **22** afforded acid **23**, which was subsequently coupled with pyrrolidine using EDC-HOBt to give **6**. Attempts to form **6** by converting **23** to the corresponding acid chloride and treating with pyrrolidine, as described for the indoles **5** and **7**–**14** above, were unsuccessful.

**Figure 3.**
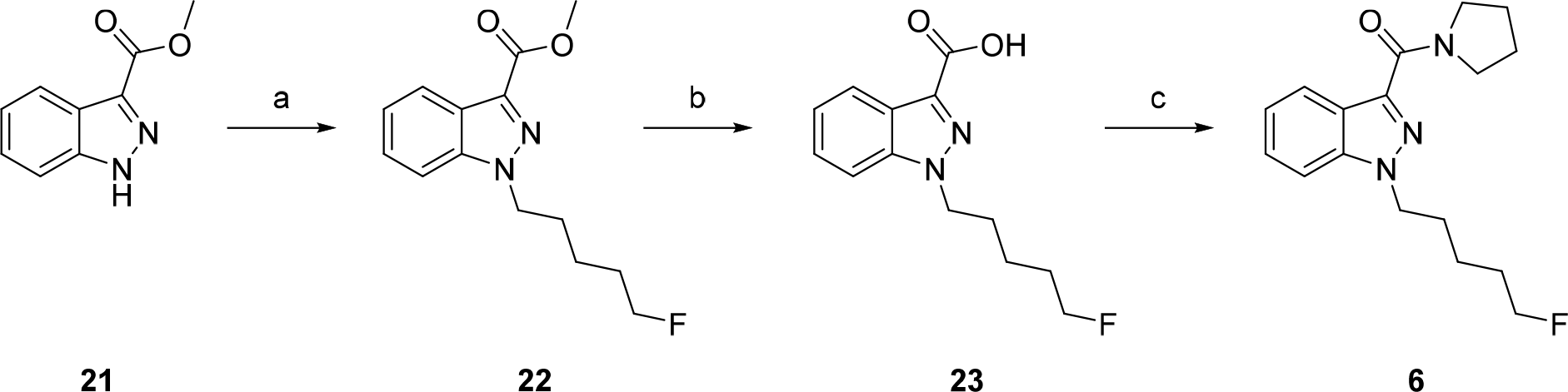
Reagents and conditions: (a) KO^t^Bu, F(CH_2_)_5_Br, THF, 0 °C-rt, 14 h, 81%; (b) 1 M aq. NaOH, MeOH, rt, 16 h, 97%; (c) EDC·HCl, HOBt·H_2_O, pyrrolidine, Et_3_N, DMF, rt, 13 h, 90%.

5F-PY-PICA, 5F-PY-PINACA, and analogues **7**–**14** were analyzed using melting point range determinations, ^1^H and ^13^C nuclear magnetic resonance (NMR) spectroscopy, and liquid chromatography–quadrupole time-of-flight–mass spectrometry (LC-QTOF-MS). Full details of these analyses are provided in the experimental section. The mass spectra and fragment assignments for 5F-PY-PICA and 5F-PY-PINACA from LC–QTOF–MS/MS analysis are shown in Figures 4a and 4b, respectively. The QTOF mass fragmentation profiles of 5F-PY-PICA and 5F-PY-PINACA were similar, with molecular ions and fragments arising from scission of the amide C-N bond observed in each case. The base peak for 5F-PY-PICA was the molecular ion (*m/z* 303.1864, 100%), and an ion consistent with loss of pyrrolidine was next most abundant (*m/z* 232.1127, 55%). The reverse was observed for 5F-PY-PINACA, with the acylium ion resulting from amide C-N bond scission occurring as the base peak (*m/z* 233.1085, 100%), and the molecular ion as the next most abundant species (*m/z* 304.1799, 54%). For 5F-PY-PICA, the only other abundant fragment (>10%) was *m/z* 98.0595 (37%), likely formed from cleavage of the indole C3-amide bond. Other minor peaks were consistent with amide cleavage and concomitant indole N1 dealkylation (*m/z* 144.0426, 8%) or defluorination (*m/z* 283.1804, 3%).

**Figure 4.**
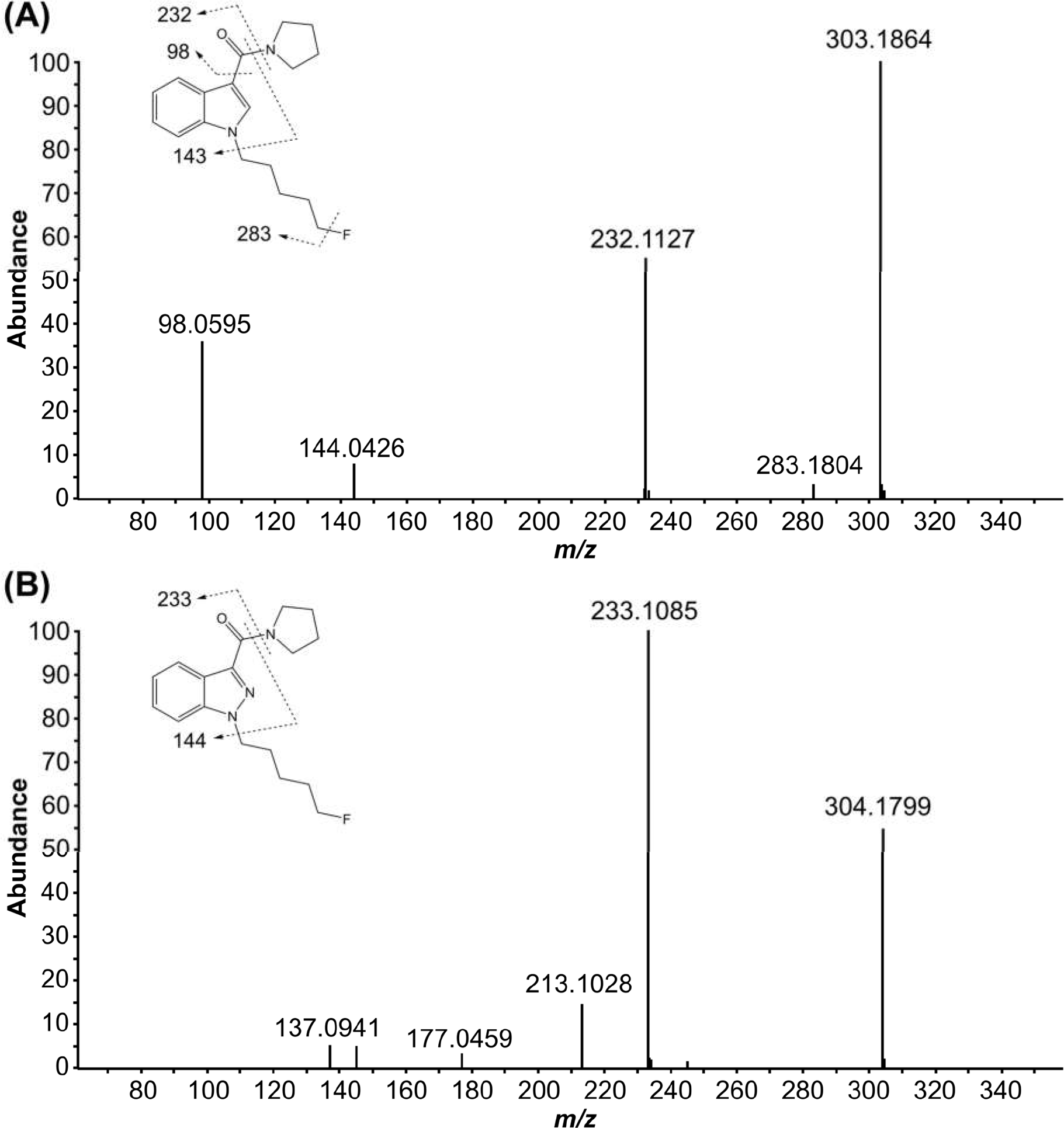
LC-QTOF-MS/MS mass spectra fragmentation patterns for (a) 5F-PY-PICA and (b) 5F-PY-PINACA.

5F-PY-PICA, 5F-PY-PINACA, and analogues **7**–**14** were screened in competitive radioligand binding assays and fluorescence-based functional assays against CB_1_ and CB_2_ receptors (Table 1).

**Table 1.**
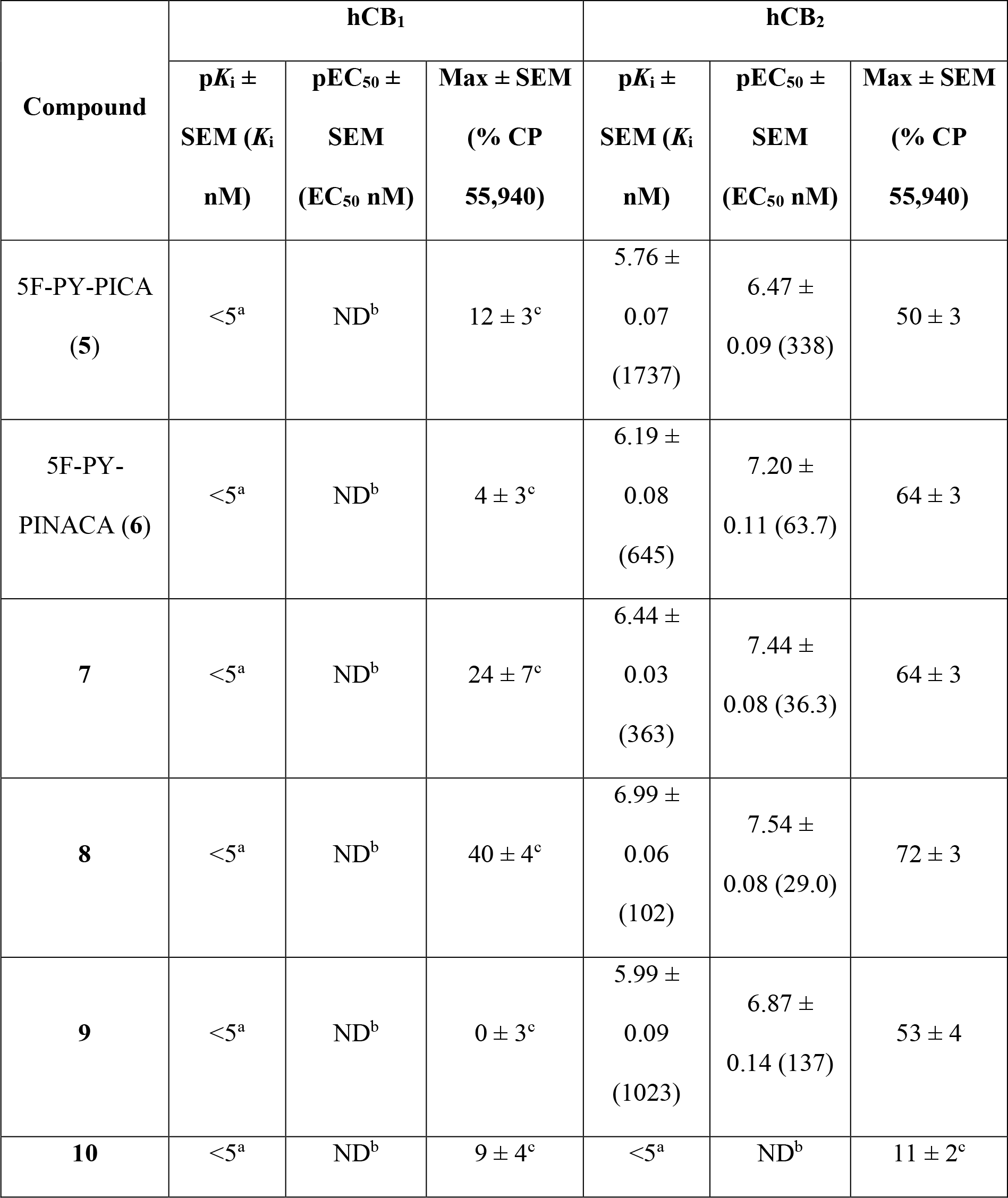
Binding affinities and functional activities of compounds **5-14** at hCB_1_ and hCB_2_ receptors.^a,b,c^

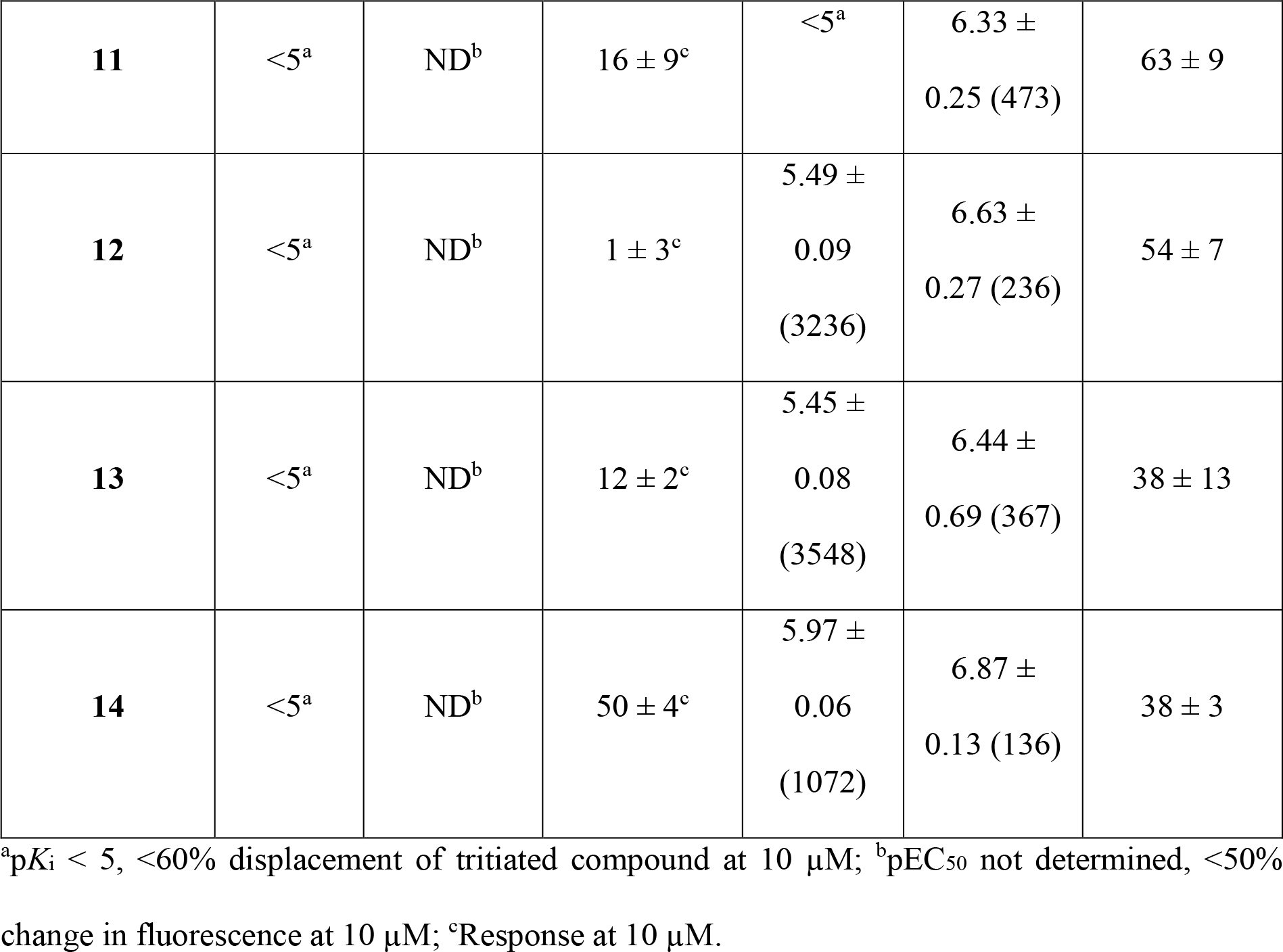

Surprisingly, despite the detection of 5F-PY-PICA and 5F-PY-PINACA as putative NPS due to their presence in drug markets, neither compound exhibited high affinity for CB_1_ (or CB_2_) receptors, with less than 60% displacement of tritiated ligand for each compound at each receptor. Additionally, analogues **7**–**14** also demonstrated minimal displacement of [^3^H]SR141716A from CB_1_, however, 5F-PY-PICA and 5F-PY-PINACA showed moderate affinities for CB_2_ (*K*_i_ = 1.73 μM and 0.65 μM, respectively). There were also clear structure-affinity trends for analogues **7**– **14**, with expansion of the pyrrolidine ring of 5F-PY-PICA by one (piperidine **7**, *K*_i_ = 363 nM) or two carbon atoms (azepane **8**, *K*_i_ = 102 nM) increasing CB_2_ affinity.

Symmetrical scission of the pyrrolidine ring of 5F-PY-PICA to the *N*,*N*-diethyl analogue also improved CB_2_ affinity (**9**, *K*_i_ = 1.02 μM), while further truncation did not (*N*,*N*-dimethyl, **10**, *K*_i_ > 10 μM). Taken together, the data suggest that 1-(5-fluoropentyl)-1*H*-indole-3-carboxamide is a suitable scaffold for the development of CB_2_ agonists, subject to sufficient steric or lipophilic contribution from pendant amide substituents.

Replacing the 5-fluoropentyl group of 5F-PY-PICA with other pendant motifs commonly seen in SCRAs (**12**–**14**) had little effect on CB_2_ binding, conferring micromolar activity in all cases (*K*_i_ = 1.07-3.55 μM) except **11** (*K*_i_ > 10 μM).

CP55,940 produced a hyperpolarization of AtT20-CB_1_ cells with a pEC_50_ of 7.76 ± 0.4, and a maximum decrease in fluorescence of 33 ± 1 %. At 10 μM, only **14** (50 ± 4%) and **8** (40 ± 4%) produced responses that were a substantial fraction of that produced by a maximally effective concentration of CP55,940 (1 μM). CP55,940 hyperpolarized AtT20-CB_2_ cells with a pEC_50_ of 7.45 ± 0.03, and a maximum decrease in fluorescence of 32 ± 1%. Several ligands produced substantial hyperpolarizations of AtT20-CB_2_ cells, but none had a maximum effect greater than 75% of CP55,940 at 1 μM. The largest activation of CB_2_ was induced by **8** (72 ± 3%).

The in vitro pharmacological profiles of 5F-PY-PICA and 5F-PY-PINACA indicate that these putative SCRAs have very low affinity and efficacy at CB_1_ receptors, and are unlikely to be psychoactive in humans. Similar to many SCRA NPS, both 5F-PY-PICA and 5F-PY-PINACA had substantial activity at CB_2_ receptors, but this is unlikely to mediate any discernable effects on mood in people.

To further characterize these compounds, and determine if they could be metabolized into active cannabinoids, we examined the effects of 5F-PY-PICA and 5F-PY-PINACA in mice using biotelemetry. The pharmacological effects of numerous SCRAs have been described in mice and rats, and include robust hypothermia, bradycardia, and hypolocomotion mediated by central CB_1_ agonist activity.^47–51^ As anticipated from the *in vitro* pharmacological data, neither 5F-PY-PICA nor 5F-PY-PINACA produced physiological effects consistent with central CB_1_ activity, and hypothermic effects were not observed in mice at doses up to 10 mg/kg (Figure 5). Many SCRAs, including JWH-018, AB-FUBINACA, MDMB-FUBINACA, and 5F-CUMYL-P7AICA, elicit hypothermic effects at doses of 1 mg/kg or below in this rodent model via a CB_1_ receptor-mediated mechanism.^13,43,52^ Therefore, it can be concluded that neither 5F-PY-PICA nor 5F-PY-PINACA exhibit the *in vitro* and *in vivo* cannabimimetic profiles shared by SCRA NPS. This does not preclude, of course, the possibility of psychoactive effects generated through non-cannabinoid mechanisms.

**Figure 5.**
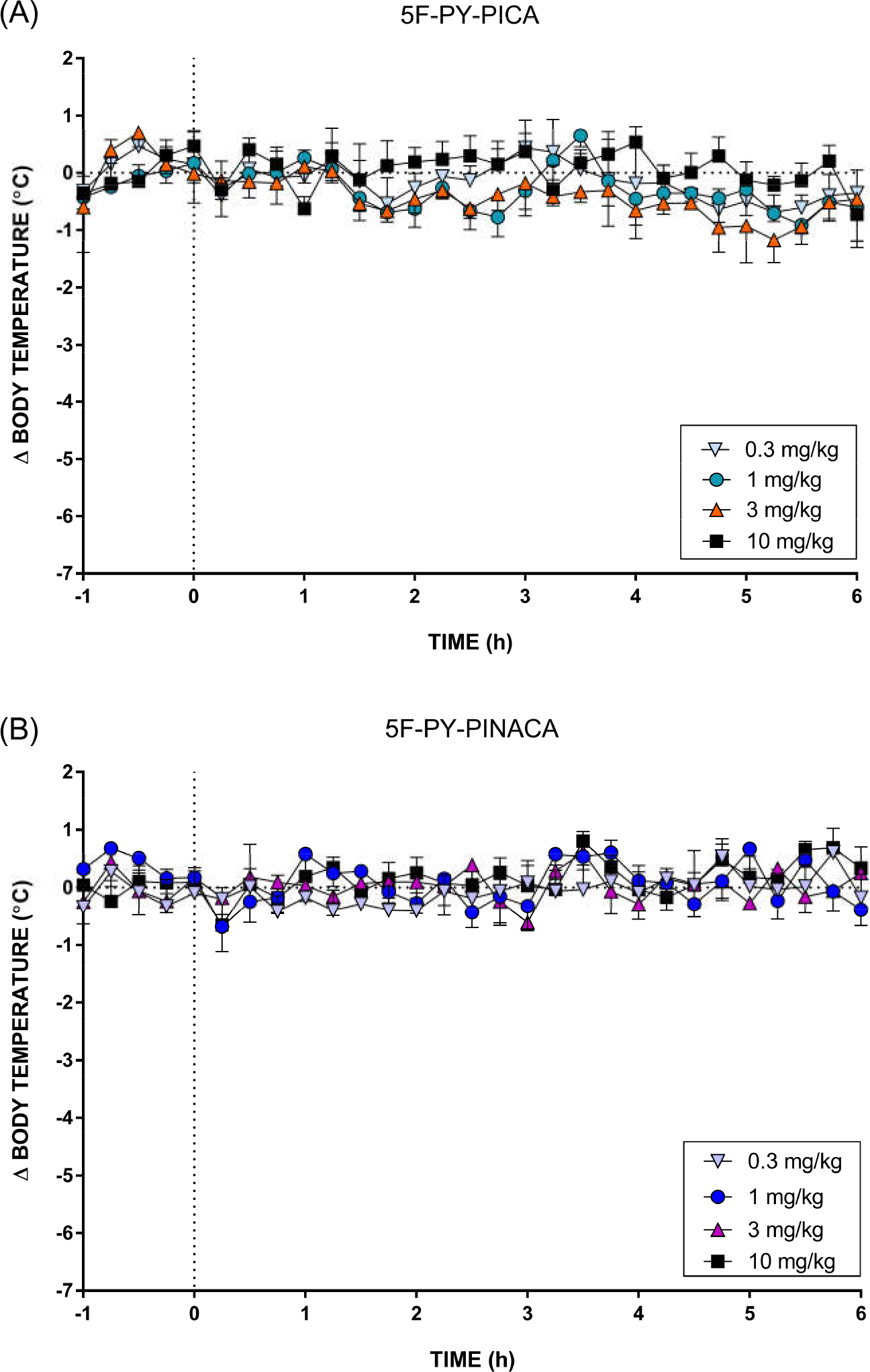
Change in body temperature following intraperitoneal injection of (A) 5F-PY-PICA and (B) 5F-PY-PINACA relative to a baseline vehicle injection. The vertical dashed line denotes time of injection, and each data point represents the mean (±SEM) change in body temperature of four mice. No hypothermic effects were observed at doses up to 10 mg/kg.

## Conclusion

Despite their detection in drug markets in 2015, 5F-PY-PICA, 5F-PY-PINACA, and several analogues exhibited low affinity and efficacy at CB_1_ and CB_2_ receptors *in vitro*. Moreover, 5F-PY-PICA and 5F-PY-PINACA failed to elicit the hypothermic, bradycardic, and hypolocomotive effects potently induced by other SCRA NPS. Taken together, the *in vitro* and *in vivo* profiles of 5F-PY-PICA and 5F-PY-PINACA cast doubt on their classification as SCRAs, although NPS designation by non-cannabinoid mechanisms cannot be excluded.

## Acknowledgement

Iain S. McGregor and Mark Connor gratefully acknowledge financial support from the Australian National Health and Medical Research Council (NHMRC Project Grant 1107088).

